# Early detection of cryostorage tank failure using a weight-based monitoring system

**DOI:** 10.1101/455311

**Authors:** Zahava P. Michaelson, Sai T. Bondalapati, Selma Amrane, Robert W. Prosser, Daniel M. Hill, Pallavi Gaur, Matt Recio, David E. Travassos, Mikaela D. Wolfkamp, Sasha Sadowy, Colin Thomas, Eric J. Forman, Zev Williams

## Abstract

**Object:** To study the ability of custom-built, web-enabled scales to monitor liquid nitrogen (LN_2_) levels in cryostorage dewars.

**Design:** Laboratory study

**Setting:** A large academic fertility center in New York City.

**Interventions:** Cryostorage dewars were placed on top of the custom-engineered scales with continuous real-time monitoring, and weight and temperature data were recorded in the setting of slow, medium, and fast rate-loss of LN_2_ designed to mimic models of tank failures.

**Main Outcome Measures:** Weights were continuously monitored and recorded, with a calculated alarm trigger set at 10% weight loss. Temperature within the tanks was simultaneously monitored with probes placed near the top of the tanks, with calculated alarms using a −185 °C as the threshold. For the “slow rate-loss” simulations, tanks were left intact and closed in usual operating conditions, and LN_2_ was allowed to evaporate at the normal rate. For the “medium rate-loss” simulation, the foam core of the tank neck was removed and the insulating vacuum was eliminated by making a 1/16 inch hole in the outer tank wall. For the “fast rate-loss” simulation, a 1/16” hole was made through the outer tank wall and LN_2_ was released at a rate of 0.15 L/second. All simulations were performed in duplicate.

**Results:** With an intact and normally functioning tank, a 10% loss in LN_2_ occurred in 4.2-4.9 days. Warming to −185 °C occurred in 37.8 - 43.7 days, over 30 days after the weight-based alarm was triggered. Full evaporation of LN_2_required 36.8 days. For the medium rate-loss simulation, a 10% loss in LN_2_ occurred in 0.8 h. Warming to −185 °C occurred in 3.7 - 4.8 hours, approximately 3 hours after the weight-based alarm was triggered. For the fast rate-loss simulation, a 10% weight loss occurred within 15 seconds and tanks were completely depleted in under 3 minutes. Tank temperatures began to rise immediately and at a relatively constant rate of 43.9 °C/hour and 51.6 °C/hour. Temperature alarms would have sounded within 0.37 and 0.06 hours after the breech.

**Conclusions:** This study demonstrates that a weight-based, automated alarm system can detect tank failures prior to a temperature-based alarm system, in some cases over a month in advance. In combination with existing safety mechanisms such as temperature probes, a weight-based monitoring system could serve as a redundant safety mechanism for added protection of cryopreserved reproductive tissues.

## Introduction

Safeguarding cryopreserved embryos and gametes is one of the critical responsibilities of fertility centers and tissue banks. Cryopreservation in liquid nitrogen (LN_2_) preserves the viability of biological materials by halting their molecular processes (1), enabling their long-term storage (2-4). Failure to maintain adequate LN_2_ within a storage tank can result in stored samples thawing and, ultimately, in their total loss. In the case of stored embryos or eggs, such a loss can be catastrophic for affected patients, their partners and the fertility center (5). Thus, it is of utmost importance to develop and perfect methods to prevent the depletion of LN_2_ from cryopreservation tanks.

Cryopreserved samples are typically stored in cryostorage tanks that utilize supercooled LN_2_ to maintain a temperature of −196 °C (6). Tanks hold ~30-75L of LN_2_ within double-walled, insulated metal containers with vacuum chambers between their two walls. Incidents of unintentional thaws of cryopreserved tissues are usually caused by one of three mechanisms (7): (A) slow rate-loss: in the case of “tank abandonment,” the cryopreservation tank functions normally and LN_2_ evaporates at the expected rate, but the tank is not refilled in a timely manner, allowing LN_2_ to be depleted and the temperature to rise; (B) medium rate-loss: a breach to the insulation of the tank, due to either damage to the vacuum seal or someone forgetting to replace the tank’s foam insulation, leads to a faster than normal rate of LN_2_ evaporation and the supply being depleted prior to the next scheduled refilling; or (C) fast rate-loss: a catastrophic tank failure causes direct leakage of LN_2_.

Existing methods to prevent unintentional thawing of cryopreserved tissues consist of a combination of manual checks and automated monitoring systems (8). The most common form of automated monitoring involves using temperature probes, which are placed within tanks and communicate wirelessly to a central monitoring station. When temperatures rise, the probes send an audible alert and, typically, automatically call a predetermined phone number. While this system is helpful, it has some inherent limitations. Firstly, temperatures only start to rise once the LN_2_ is nearly depleted. There is, in turn, a very limited timeframe for which the LN_2_ can be replaced before the specimens are completely lost. Secondly, since the sensors note a rise in temperature, the lowest possible storage temperature will have, by definition, been passed before the alarm is sounded. Thirdly, this method requires that sensitive electronic equipment be placed within the harsh conditions of a cryopreservation tank, thereby limiting the lifespan of the probes and increasing the cost for monitoring. Finally, because current monitors require that probes be placed within the LN_2_, the introduction of a second probe for additional monitoring of samples is cumbersome, and represents a replicative rather than truly redundant method. There is, as such, a need for a monitoring method that is redundant to the temperature probe, is positioned externally to the tank, and provides early detection of impending tank failure to safeguard the integrity of cryopreserved tissues.

We hypothesized that continuous observation of tank weight could be a simple, safe, low-cost method for real-time, automated monitoring of cryostorage tanks. In contrast to temperature alarms, weight-based alarms can be calibrated to sound when a given proportion of LN_2_ is lost. An alert would thus be triggered *before* the LN_2_ supply is completely depleted and the tank’s temperature rises, giving staff more time to refill the tank or transfer samples to a different tank. The scale would also be external to the tank, thereby obviating the need for electronics that could withstand the supercooled environment within the tanks. Finally, a weight-based alarm system could provide a redundancy to the existing safety mechanisms.

Here we tested the ability of custom-built, web-enabled scales to monitor cryostorage dewars in slow, medium and fast rate-loss tank failure simulations and demonstrate that a weight-based system provides an early-warning and redundant safeguard for cryopreserved specimens.

### Materials and Methods

For all simulations, custom-engineered and manufactured scales were used to measure and record tank weight. The scales had a 400 lb capacity and data output capabilities, and were attached to a terminal that was connected via Ethernet to a central computer monitoring station. TCP Wedge with custom scripts was used for collection of six fields of data from the networked device. A programmable alarm system was used to record real-time weight information (Figure 1). Alarms were set to trigger after a 10% weight loss. All experiments were performed with 22.4L MDD 93/42/EEC compliant passive cryostorage tanks (MVE xc 22/5; Chart Biomedical) containing empty canisters and no patient samples. Tanks were placed on the scales and the scales were tared. The tanks were then filled with LN_2_, and topped-off until temperature reached equilibrium and LN_2_ boiling ceased. Temperature within the tanks was simultaneously monitored with probes (Safepoint Scientific) placed near the top of the tanks with calculated alarms using a −185 °C threshold, as per standard operating practice. Ambient room temperatures were 23-25 °C.

**Figure 1:**
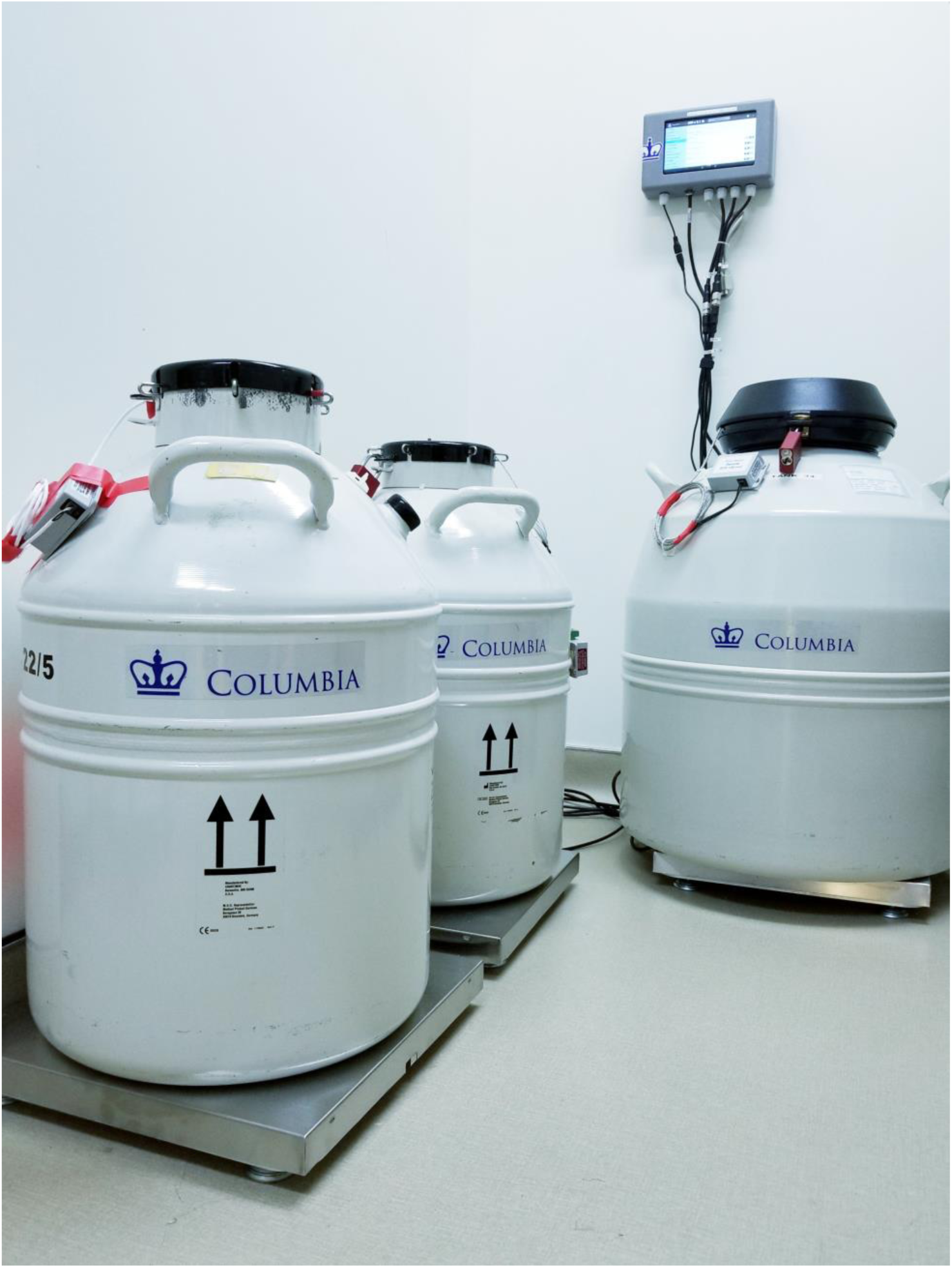
Experimental set-up. LN_2_ dewars equipped with temperature probes were placed on scales. Weights were recorded continuously in real-time via relay to an Ethernet-connected monitoring unit.

For the “slow rate-loss” simulations, tanks were left intact and closed and in usual operating conditions. LN_2_ was allowed to evaporate at the normal rate until all LN_2_ had evaporated and temperature returned to room temperature (RT). For the “medium rate-loss” simulation, the foam core of the tank neck was removed and the insulating vacuum was eliminated by the creation of a 1/16” hole in just the outer tank wall. For the “fast rate-loss” simulation, a 1/16” hole was made through the outer tank wall and LN2 was released at a rate of 0.15 L/second. All simulations were performed in duplicate.

## Results

For the slow rate-loss simulation, designed to mimic what would occur if a fully functioning tank was left unattended, two LN_2_ dewars (Tank A and Tank B) were filled to capacity with LN_2_, topped-off until temperature reached equilibrium and LN_2_ boiling ceased, and then left undisturbed while the tank weights and temperatures were monitored continuously (Figure 2, Table 1). The time required for a 10% loss in LN_2_ was 4.2 and 4.9 days for Tanks A and B, respectively (Table 1). Warming to −185 °C occurred in 37.8 and 43.7 days for Tanks A and B, respectively. Thus, the weight-based system was able to detect a 10% weight loss between 33.7 and 38.8 days before the temperature alarm threshold was reached (Figure 2A and B, Table 1). The rate of LN_2_ loss was 1.05 kg/day and 0.89 kg/day for Tank A and B, respectively. Because Tank B retained LN2 better, Tank A was used for subsequent studies to minimize the differences between weight and temperature based monitoring. Full evaporation of Tank A required 36.8 days (Figure 2A). Tank temperatures, as recorded by a temperature probe placed a few centimeters below the tank neck, remained stable until LN_2_ was at or nearly completely depleted (Figure 2B). Thereafter, the rate of warming was nearly constant and occurred at a rate of and 60.5 °C/day (Figure 2B). The rate of weight loss for the intact tank can be described by the formula W=−2.778*t* + 101.48, where W is the weight as a percentage of the full tank and *t* is time in days (R^2^= 0.9999).

**Figure 2:**
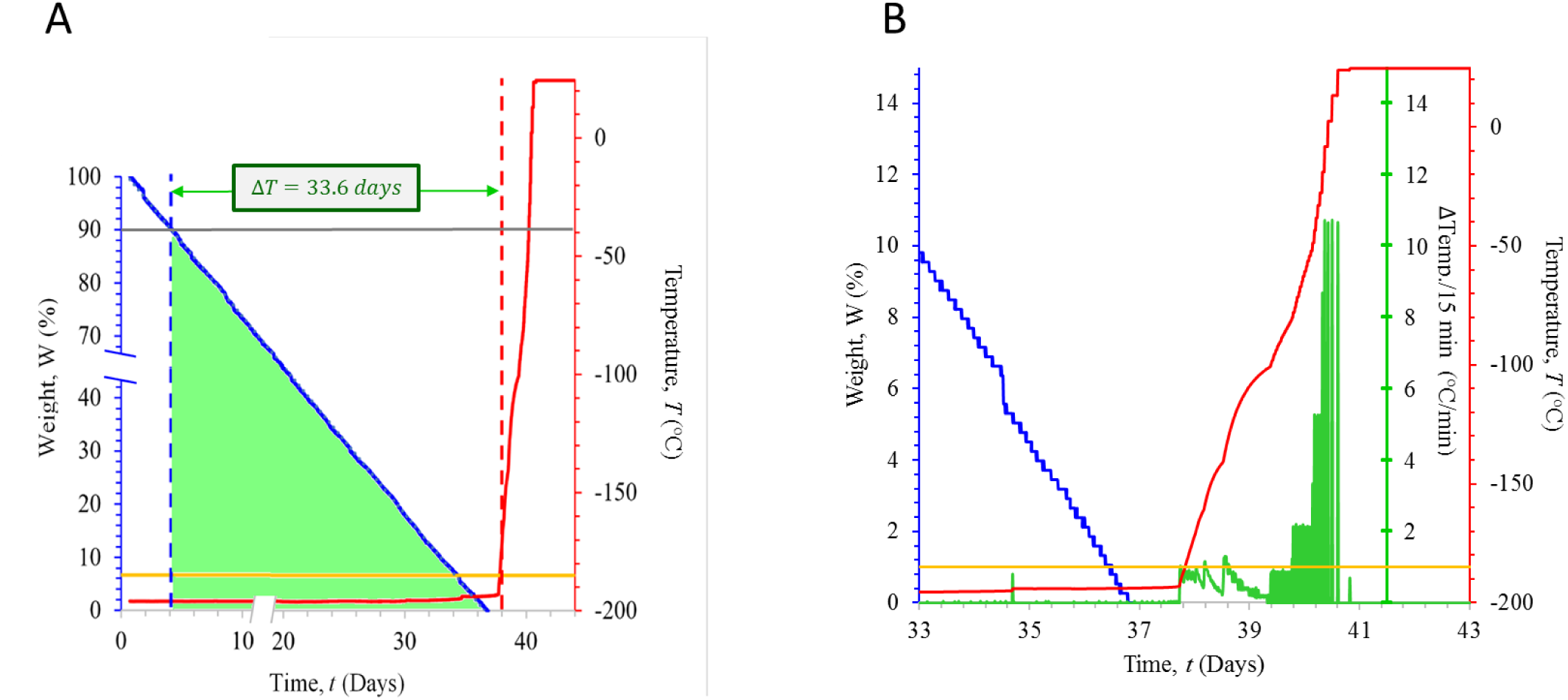
Results of the slow rate-loss trial. **(A)** The decrease in LN_2_ weight (as a percentage of starting weight; Blue line), and the corresponding increase in temperature (°C) over time (measured in days; Red line) are shown. Thresholds for detecting a 10% weight loss and a temperature of −185 °C are indicated by the grey and yellow lines, respectively. The interval between the detection of 10% weight drop and a rise in temperature to −185 °C is 33.6 days and is indicated as the distance between the dashed blue and red lines. The rate of weight loss can be described by the formula W = −2.778*t* + 101.48 where W is the LN_2_ weight (as a percentage of total) and *t* is the time (measured in days; R^2^ = 0.9999). **(B)** Expansion of the period of time between days 33 and 43 shows a slight time-interval between when the tank fully empties (blue line) and when the temperatures begin to rise (red line). The threshold for triggering the - 185 °C temperature alarm is indicated in yellow. The rate of temperature increase, shown as the change in temperature (°C) for each 15 min interval is indicated in green.

**Table 1:** Time require for a 10% drop in weight LN_2_ from a full tank, the time required for temperatures to rise to −185 °C and the interval between those two times for each of the experimental conditions.

|  | Slow-rate<br>Tank A<br>(Days) | Slow-rate<br>Tank B<br>(Days) | Medium-rate<br>Run 1<br>(Hours) | Medium-rate<br>Run 2<br>(Hours) | Fast-rate<br>Run 1<br>(Hours) | Fast-rate<br>Run 2<br>(Min) |
| --- | --- | --- | --- | --- | --- | --- |
| Time to 10% ↓wt | 4.2 | 4.9 | 0.8 | 0.8 | NA | NA |
| Time to -185 °C | 37.8 | 43.7 | 3.7 | 4.8 | 0.37 | 0.06 |
| △ 10%↓wt and -185°C | 33.7 | 38.8 | 3.0 | 4.0 | NA | NA |

For the medium rate-loss simulation, designed to mimic what would occur if the insulating capacity of a tank were compromised, the tank’s vacuum seal was breached and insulating foam in the tank neck was removed. Dewars were then filled to capacity with LN_2_, topped-off until temperature reached equilibrium and LN_2_ boiling ceased and then left alone while the tank weight and temperature was monitored continuously. The time required for a 10% loss in LN_2_ was 0.8 hours for runs 1 and 2 (Table 1, Figure 3A). Warming to −185 °C occurred in 3.7 and 4.8 hours for runs 1 and 2, respectively (Table 1). Thus, the weight-based system was able to detect a 10% weight loss between 3.0 and 4.0 hours before the temperature alarm threshold was reached (Figure 3A, Table 1). The rate of LN_2_ loss was significantly higher than in the trials using the uncompromised tanks and was fastest for the loss of the initial 50% of LN_2_: the first 50% of volume was lost in 4.3 and 4.7 hours for runs 1 and 2, respectively, while the remaining 50% of tank volume and was lost in 7.5 and 7.8 hours for runs 1 and 2, respectively Figure 3A and Supplementary Figure 1). In contrast to the results of the uncompromised tank study, temperatures in the compromised tanks began rising much sooner—just about one hour after 10% of LN_2_ volume was lost. The rate of temperature rise was bi-phasic with a slower rate of warming (3 °C/hour and 2.3 °C/hour for run 1 and 2, respectively) while LN_2_ was present and a much faster rate of warming (48 °C/hour and 58.3 °C/hour for run 1 and 2, respectively) after LN_2_ was depleted (Figure 3A and Supplementary Figure 1). The rate of weight loss for the breeched tank can be approximated by the formula W= 0.4447*t* ^2^- 13.661*t* + 100.07 where W is the weight as a percentage of the full tank and *t* is time in hours. (R^2^=0.9997, calculated from Run 1)

**Figure 3:**
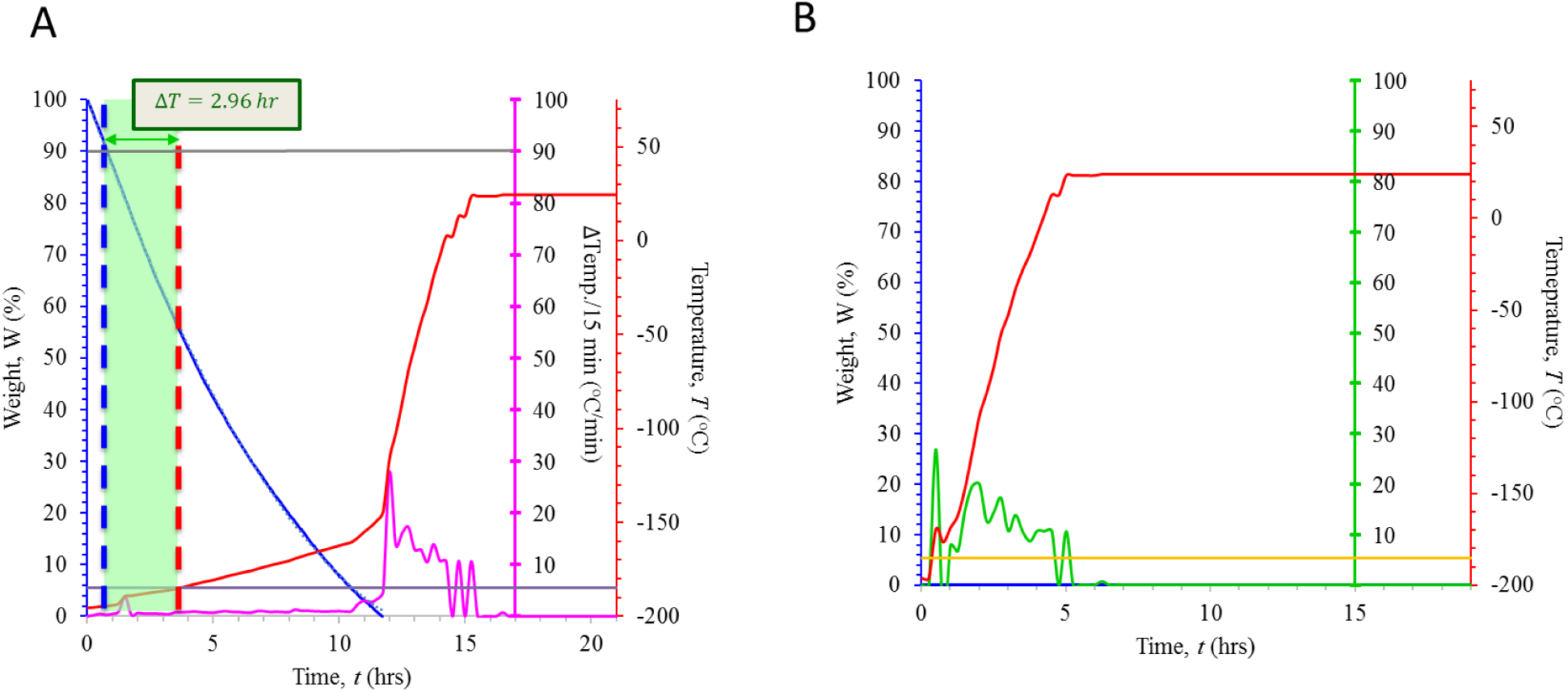
(A) Results of the medium rate-loss trial. The decrease in LN_2_ weight (as a percentage of starting weight; Blue line), and the corresponding increase in temperature (°C) over time (measured in hours; Red line) are shown. Thresholds for detecting a 10% weight loss and rise in temperature to −185 °C are indicated by the grey and purple lines, respectively. The interval between the detection of 10% weight drop and a rise in temperature to −185 °C was approximately 3 hours and is indicated as the distance between the dashed blue and red lines. The rate of weight loss can be described by the formula W = 0.4447*t* ^2^ - 13.661*t* + 100.07 (where W represents weight, measured as a percentage of total and *t* represents time, measured in hours; R^2^ = 0.9997). The rate of change of temperature, measured in 15 minute intervals and recorded as °C/min is shown in magenta. **(B) Results of the rapid rate-loss trial.** Thresholds for detecting a rise in temperature to −185 °C are indicated by the yellow lines. The rate of change of temperature, measured in 15 minute intervals and recorded as °C/min is shown in green.

For the fast rate-loss simulation, designed to mimic what would occur if a catastrophic tank breech were to cause LN_2_ to rapidly spill from the tanks, the tanks’ vacuum seal was breached. Dewars were then filled to capacity with LN_2_, topped-off until the temperature reached equilibrium and LN_2_ boiling ceased, and the LN_2_ was evacuated at a rate of 0.15 L/second until empty. The tanks were left alone while their temperature and weight were monitored continuously (Figure 3B, Table 1, Supplementary Figure 2). Thus, a 10% weight loss occurred within 15 seconds and tanks were completely depleted in under 3 minutes. As shown in Figure 3B, tank temperatures began to rise immediately and at a relatively constant rate of 43.9 °C/hour and 51.6 °C/hour. Temperature alarms would have sounded within 0.37 and 0.06 hours after the breech (Figure 3B, Table 1, Supplementary Figure 2).

## Discussion

Existing methods to protect cryopreserved reproductive tissue are reliable and failures are extremely rare (5). However, as recent cases have highlighted, failures are still possible and thus there is a need for additional safeguards.

This study demonstrates that a weight-based, automated alarm system can detect tank failures prior to a temperature-based alarm system. The time interval between the weight-based and temperature-based alarms varies depending on the threshold set for each and the rapidity and nature of the loss. Using standard temperature alarm settings and a 10% weight-loss threshold, the weight-based alarm detected anomalies as far as 33 days in advance of a temperature rise. In cases where there was a higher rate of LN_2_ loss, the interval was smaller.

When the insulating capacity of the tanks were left intact, temperatures remained low until several days after LN_2_ was completely depleted whereas, when the insulating capacity was breached, temperatures began rising much sooner. This is likely because, with the insulation intact, supercooled nitrogen vapor was able to maintain low tank temperatures, but this ability failed without the vacuum layer. This is of practical implication because the normal buffer time available to replenish LN_2_ is drastically shortened in cases of the loss of insulating capacity.

It should be noted that the anticipated application of this technology is not as a replacement for existing safety monitoring systems, including temperature, but as an additional and redundant safety mechanism. Since existing tanks and their probes can be placed directly on scales, this technology can be readily adopted. For centers that place tanks on wheeled dollies, the scale can be designed to fit onto the dolly and beneath the dewar, or integrated into the dolly itself. In cases of cryopreservation of less critical tissues (e.g. animal cell lines), a weight-based alarm system may be a cost-effective alternative to temperature-based alarms. Given the ability to accurately record the rate of loss of LN_2_, the scales can also be used for QC testing of tanks, including new tanks, to confirm that they maintain LN_2_ as specified.

There were several limitations to this study. The first is that it was conducted with a single brand and make of dewar. There is likely variation in the rate of LN_2_ loss and temperature rise depending on the particular make, model and integrity of the dewar. The time interval between when weight-based and temperature-based alarms trigger also depends on where within the tank the temperature probe is placed. In our experiments, temperature probes were placed near the top of the tank. Placing the temperature probe near the bottom of the tank would likely result in even more LN_2_ lost before a temperature probe would detect a change. Performance of the scale-based system in real-world conditions and over time will also need to be evaluated. The threshold for triggering the temperature was set at −185 °C. Published recommendations for setting the temperature threshold for triggering the temperature-based alarm vary from −185 °C and warmer (8). We chose the −185 °C in order to minimize the interval between the weight and temperature-based thresholds. Using a warmer temperature threshold would be expected to extend the interval between when the weight-based and temperature-based alarms would trigger and would also reduce the time available to respond before samples were lost. Finally, some centers use nitrogen vapor tanks for storage of cryopreserved reproductive tissue (8) and the kinetics of a failure of these tanks remains to be determined, though we would anticipate that, while weight would still change prior to a temperature rise, the interval may be shorter.

While risk can never be completely eliminated, our hope is that by adding additional safeguards to the existing manual and automated safety mechanisms, cryopreservation can become even safer and tank failures can become as impossible as possible.

## Acknowledgements

The authors thank the members of Columbia University Fertility Center for their assistance in conducting the trial. ZW is partially supported through funding from the National Institutes of Health (grants HD068546 and U19CA179564). ZW is listed as an inventor on patent applications filed by Columbia University relating to this system.

---

**Supplementary Figure 1:**
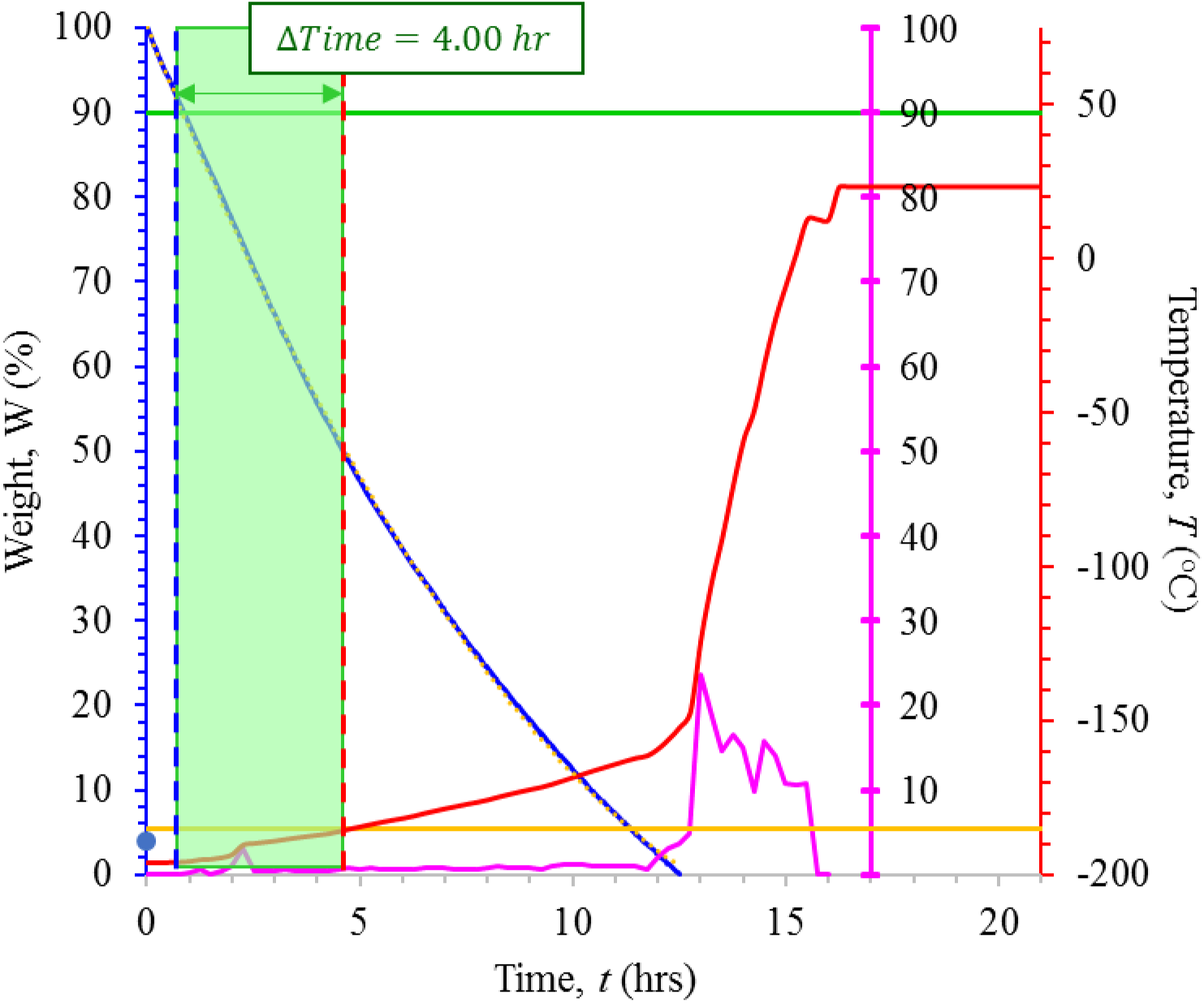
Results of the repeat run of the medium rate-loss trial. The decrease in LN_2_ weight (as a percentage of starting weight; Blue line), and the corresponding increase in temperature (°C) over time (measured in days; Red line) are shown. Thresholds for detecting a 10% weight loss and rise in temperature to −185 °C are indicated by the grey and yellow lines, respectively. The interval between the detection of 10% weight drop and a rise in temperature to −185 °C was approximately 4 hours and is indicated as the distance between the dashed blue and red lines. The rate of weight loss can be described by the formula W = *0.3565t* ^2^ - 12.383*t* + 100.09 (where W represents weight, measured as a percentage of total and *t* represents time, measured in hours; R^2^ = 0.9998). The rate of change of temperature, measured in 15 minute intervals and recorded as °C/min is shown in magenta

**Supplementary Figure 2:**
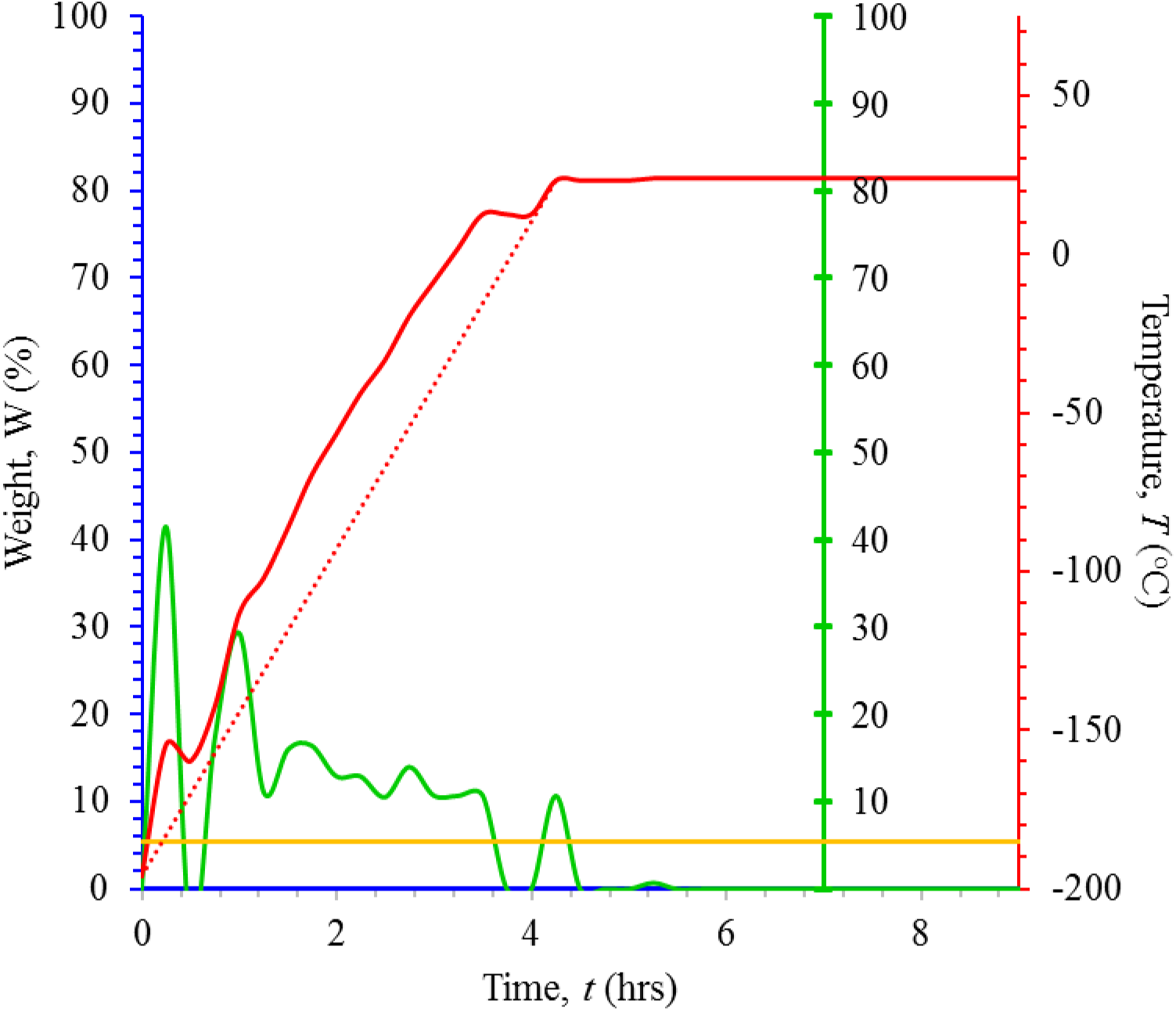
Results of the rapid rate-loss trial. Thresholds for detecting a rise in temperature to −185 °C are indicated by the yellow lines. The rate of change of temperature, measured in 15 minute intervals and recorded as °C/min is shown in green.

